# Cross-species incompatibility between a DNA satellite and the Drosophila Spartan homolog poisons germline genome integrity

**DOI:** 10.1101/2021.08.13.455988

**Authors:** Cara L. Brand, Mia T. Levine

**Affiliations:** Department of Biology and Epigenetics Institute, University of Pennsylvania, Philadelphia, PA

**Keywords:** DNA satellite, coevolution, maternal haploid, 359, Spartan, Topoisomerase II

## Abstract

Satellite DNA spans megabases of eukaryotic sequence and evolves rapidly. Paradoxically, satellite-rich genomic regions mediate strictly conserved, essential processes like chromosome segregation and nuclear structure. A leading resolution to this paradox posits that satellite DNA and satellite-associated chromosomal proteins coevolve to preserve these essential functions. We experimentally test this model of intra-genomic coevolution by conducting the first evolution-guided manipulation of both chromosomal protein and DNA satellite. The 359bp satellite spans an 11Mb array in *D. melanogaster* that is absent from its sister species, *D. simulans*. This species-specific DNA satellite colocalizes with the adaptively evolving, ovary-enriched protein, Maternal Haploid (MH)–the Drosophila homolog of Spartan. To determine if MH and 359 coevolve, we swapped the *D. simulans* version of MH (“MH[sim]”) into *D. melanogaster*. MH[sim] triggers ovarian cell death, reduced ovary size, and loss of mature eggs. Surprisingly, the *D. melanogaster mh* null mutant has no such ovary phenotypes, suggesting that MH[sim] is toxic in a *D. melanogaster* background. Using both cell biology and genetics, we discovered that MH[sim] poisons oogenesis through a DNA damage pathway. Remarkably, deleting the *D. melanogaster*-specific 359 satellite array completely restores *mh[sim]* germline genome integrity and fertility, consistent with a history of coevolution between these two fast-evolving loci. Germline genome integrity and fertility are also restored by overexpressing Topoisomerase II (Top2), suggesting that MH[sim] interferes with Top2-mediated processing of 359. The observed 359-MH[sim] cross-species incompatibility supports a model under which ostensibly inert repetitive DNA and essential chromosomal proteins must coevolve to preserve germline genome integrity.

## RESULTS AND DISCUSSION

DNA satellite-enriched genomic regions evolve rapidly and yet support strictly conserved nuclear functions.^1–9^ A classic resolution to this paradox posits that DNA satellite-associated proteins evolve adaptively to mitigate deleterious proliferation of DNA satellite sequence variants.^10^ Repeated bouts of DNA satellite evolution and chromosomal protein adaptation result in exquisitely coevolved satellites and satellite-associated proteins. This model of coevolution predicts pervasive incompatibilities between satellite DNA and chromosomal proteins from closely related species: adaptively evolving chromosomal proteins from one species should fail to package or process DNA satellites from another.^10–12^

Evidence for this coevolution model has emerged from engineering “evolutionary mismatches” between the adaptively evolving chromosomal protein(s) of one species and the DNA satellite landscape of a close relative. Under one approach, a diverged chromosomal protein is introduced into a closely related species, generating an evolutionary mismatch between the manipulated protein and one or more DNA satellites.^12–15^ Consistent with disrupted DNA satellite:chromosomal protein coevolution, the naïve protein typically perturbs a satellite-mediated function, such as chromosome segregation or nuclear organization.^12,14,15^ In these cases, however, the incompatible DNA satellites are unknown. A second approach crosses sister species to generate evolutionary mismatches between chromosomal proteins and DNA satellites in hybrid progeny. Consistent with disrupted DNA satellite:chromosomal protein coevolution, interspecies hybrid inviability has been linked to satellite-rich genomic loci.^16,17^ In these systems, however, the incompatible chromosomal proteins are unknown. To date, there are no cases of experimental identification of both chromosomal protein and satellite engaged in coevolution.

To experimentally probe *both sides* of the coevolution model, we searched for a rapidly evolving DNA satellite associated with an adaptively evolving chromosomal protein. In *Drosophila melanogaster*, the 359bp satellite spans an 11Mb array at the base of the *X* chromosome.^18,19^ Close relatives of *D. melanogaster*, including *D. simulans* and *D. erecta*, lack this *X*-linked satellite array.^20^ Instead, these species have shorter arrays of “359-like” sequence dispersed throughout heterochromatin and euchromatin.^9,21,22^ Such extreme lineage-restriction to *D. melanogaster* makes this DNA satellite array an ideal locus for testing the coevolution model. On the protein side, we identified from the literature Maternal Haploid (MH), an ovary-enriched protein that is maternally provisioned to the embryo and colocalizes with the 359 satellite.^23,24^ Embryos of *mh* null mothers suffer paternal chromosome mis-segregation at the very first mitosis, suggesting that the maternally-provisioned MH prepares the otherwise inert, sperm-deposited paternal chromosomes for participation in embryonic mitosis.^23,24^ Most of these embryos arrest around the first mitotic division. A smaller fraction develop beyond the first division, cycling only the “maternal haploid” complement of chromosomes until arrest prior to hatching.^23–25^

If the *D. melanogaster*-specific 359 proliferation triggered *mh* to innovate, we should detect evidence of positive selection at *mh* between *D. melanogaster* and *D. simulans*. To determine if *mh* evolves adaptively, we conducted a McDonald-Kreitman test^26^ using polymorphism within *D. melanogaster* and *D. simulans* populations and divergence between *D. melanogaster* and *D. simulans* (2.5 million years diverged^27^). This comparison revealed an excess of nonsynonymous fixations, consistent with a history of adaptive evolution (Figure 1A). The dynamic evolution of the 359 satellite and adaptive evolution of a 359-associated protein, MH, raises the possibility that *mh* recurrently evolves to preserve a biological function compromised by 359 satellite proliferation.

**Figure 1.**
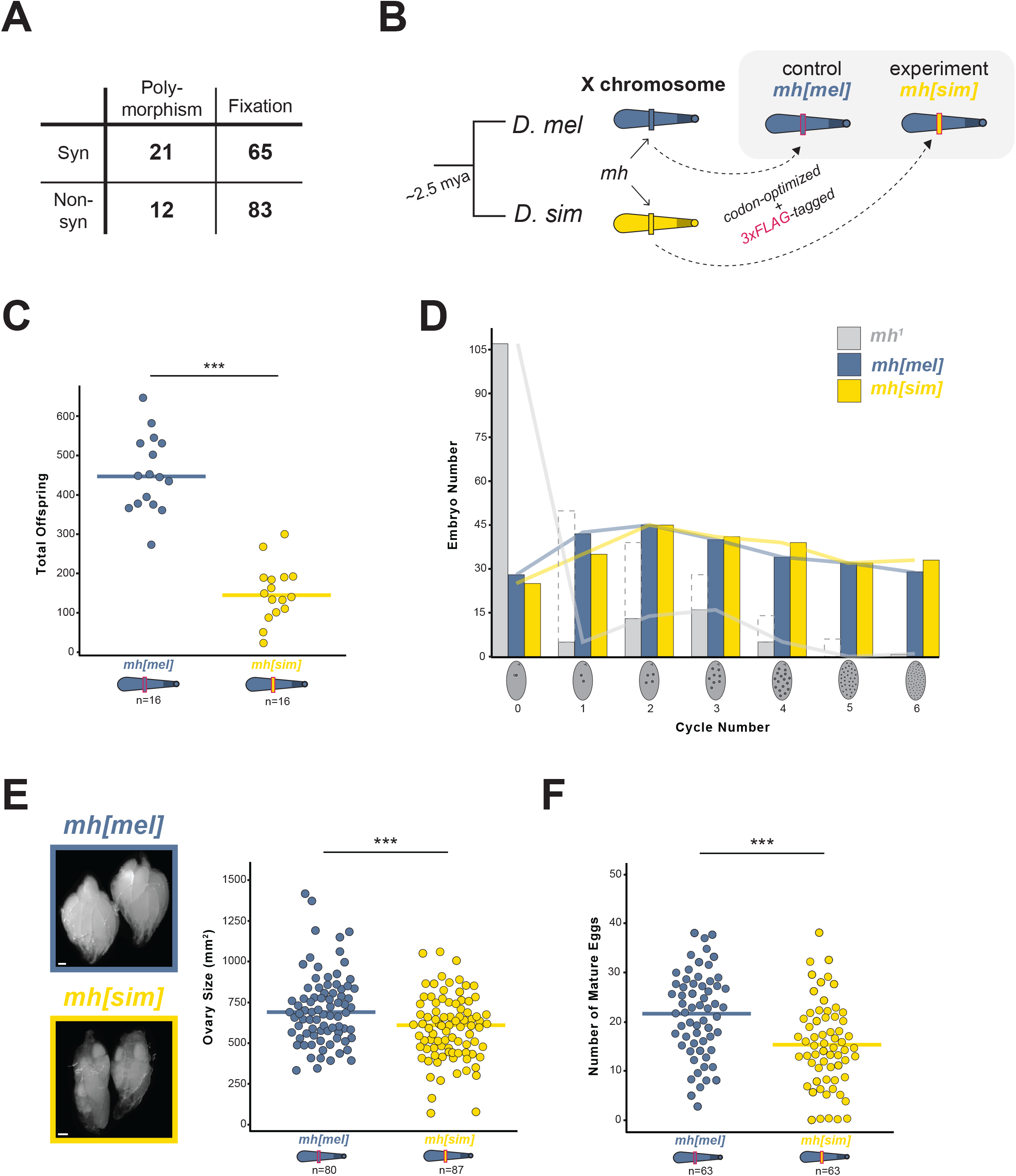
MH evolves adaptively to preserve female fertility. (A) Counts of synonymous and nonsynonymous polymorphic and fixed sites within and between *D. melanogaster* and *D. simulans; χ*^2^ test, p = 0.04. (B) Swap strategy: the *D. melanogaster* (“*D. mel*”, blue) or *D. simulans* (“*D. sim*,” yellow) *mh* coding sequence, codon-optimized for *D. melanogaster* and 3xFLAG-tagged, replaced the native *mh* gene on the *X* chromosome. (C) Total offspring from *mh[mel]* or *mh[sim]* females crossed to wildtype (*w*^*1118*^) males. (D) Frequency distribution of embryos at increasing mitotic cycle numbers collected for 70 minutes from *mh[mel]*, *mh[sim]*, and *mh*^*1*^ females. Dashed bars from *mh*^*1*^ females correspond to embryos undergoing mitotic catastrophe likely triggered at the first mitosis. Solid gray bars representing *mh*^*1*^-derived embryos are presumed to be maternal haploid. (E) Representative images and ovary size estimates from *mh[mel]* and *mh[sim]* females. (F) Number of mature eggs per ovary pair from *mh[mel]* and *mh[sim]* females. (*t*-test: “***” = p < 0.001, scale bar = 100μm)

To test the possibility of MH:359 coevolution, we first conducted an evolution-guided manipulation of *mh* to generate an “evolutionary mismatch” between protein and satellite. We used CRISPR/Cas9 to integrate into the native *mh* locus of *D. melanogaster* either a 3xFLAG-tagged *mh* coding sequence from *D. melanogaster* (our control fly, “*mh[mel]*”) or a 3xFLAG-tagged *mh* coding sequence from *D. simulans* (our experimental fly, “*mh[sim]*”, Figure 1B). Both the *D. melanogaster* and the *D. simulans* coding sequences were codon-optimized for *D. melanogaster.* We observed equivalent expression of the two transgenes (Figure S1A).

The *mh* null mutant phenotype in the early embryo motivated our prediction that an evolutionary mismatch between the *D. simulans mh* and the *D. melanogaster* 359 *X*-linked array would compromise paternal chromosome segregation during the first mitotic division. Specifically, we predicted that MH[sim] would fail to recognize and process 359, triggering mis-segregation of the paternal *X*-chromosome. This defect would result in reduced female fertility and a dearth of female progeny. We discovered that *mh[sim]* females produced significantly fewer progeny than control *mh[mel]* females (Figure 1C); however, contrary to our prediction, the progeny sex ratio did not deviate from 50/50 (Figure S1B). These data suggest that paternal 359 is *not* uniquely vulnerable to the presence of MH[sim] during the first mitotic division. Consistent with this inference, we observed that *mh[sim]* completely rescues the first mitotic division: embryos from *mh[mel]* and *mh[sim]* mothers show equivalent, normal distributions of embryonic stages from a 70-minute collection (Figure 1D). In contrast, embryos produced by *mh* null mothers typically arrest during the first division (Figure 1D). Moreover, we observed no evidence of elevated maternal haploid embryos from *mh[sim]* mothers (Figure S1C). These data suggest that *mh[sim]* does not phenocopy the *mh* null early embryonic phenotype.

To uncover an alternative source of the *mh[sim]* fertility defect, we looked at the developmental stage just before the first embryonic mitosis: oogenesis. Although *mh* is highly expressed during oogenesis, previous reports suggested that *mh* null alone yields no ovary phenotype^23,24^. We similarly detected no difference in ovary size or mature egg number of *mh* null mothers compared to heterozygous controls (Figure S1D,E). In contrast, *mh[sim]* ovaries are significantly smaller than *mh[mel]* ovaries and are depleted of the most mature egg stages (Figure 1E,F). This unexpected *mh[sim]* ovary phenotype, combined with the complete rescue of the first embryonic division by *mh[sim]*, suggests that *mh[sim]* does not behave as a loss-of-function allele. Instead, MH[sim] might be toxic.

To explore the possibility that MH[sim] is toxic, we first asked if MH[sim] localizes aberrantly in the ovary. We visualized MH[mel] and MH[sim] by staining ovaries with anti-FLAG. We discovered that MH[mel] localized primarily in the earliest stages of oogenesis (the germarium, Figure 2A,B). MH[sim] localized not only in these early stages of oogenesis but also on the “nurse cell” nuclei of later stage egg chambers and weakly on the somatic follicle cell nuclei (Figure 2A,B). The aberrant persistence of MH[sim] during oogenesis, combined with compromised *mh[sim]* ovary development, raised the possibility that MH mislocalization alone might be toxic. To test this hypothesis, we used the UAS/GAL4 system to overexpress MH[mel] in the female germline (driver nos-Gal4-VP16). In ovaries overexpressing MH[mel], we indeed observed elevated levels and aberrant persistence of the protein in later stage egg chambers (Figure S2A). Nevertheless, these females gave rise to abundant progeny (Figure S2B), suggesting that mislocalization alone cannot explain the compromised ovary development of *mh[sim]* females. In contrast, overexpression of MH[sim] in otherwise wildtype ovaries resulted in a complete absence of mature eggs (nosGAL4 > UASp-*mh[sim]*: μ = 0 eggs/ovary pair versus native > *mh[sim]*: μ = 15 eggs/ovary pair, n = 63 ovary pairs/genotype). Consequently, these females were completely sterile (Figure. S2B). These data suggest that MH[sim] – which functions normally in its native *D. simulans* genome – is toxic to oogenesis in *D. melanogaster*. This toxicity appears to be dose-dependent. Heterozygous *mh[mel]/mh[sim]* females give rise to progeny counts similar to *mh[mel]* homozygotes (Figure S2C).

**Figure 2.**
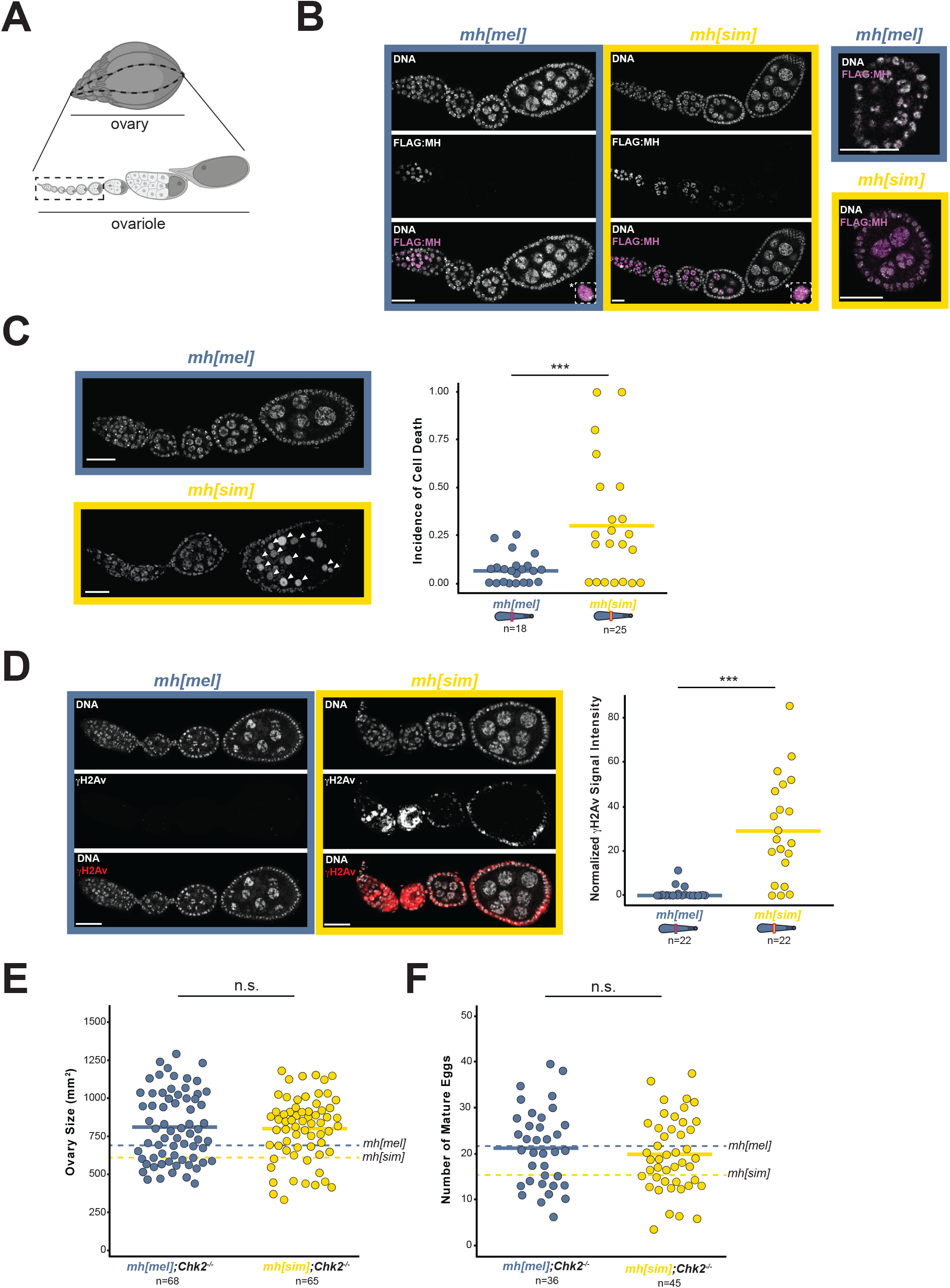
MH[sim] poisons oogenesis through a DNA damage pathway. (A) Diagram of a Drosophila ovary (above) and a single ovariole (below) with the germline stem cells in the germarium at the anterior (left) position and the mature eggs at the posterior position (right). The dashed box shows the developmental stages shown in images 2B-2D (Created using BioRender.com). (B) *mh[mel]* and *mh[sim]* ovaries stained with anti-FLAG to visualize MH localization (left). Merged images of single nuclei (*) show no MH foci on the DNA (dashed inset boxes). Merged images of enlarged egg chambers imaged under higher laser power reveal that MH[sim] but not MH[mel] localizes to the peripheral follicle cells (right). (C) Incidence of cell death captured by the fraction of ovarioles with condensed nuclei (arrowheads) in *mh[mel]* and *mh[sim]* ovaries. (D) γH2Av signal in *mh[mel]* and *mh[sim]* ovaries and the quantification of normalized fluorescent signal intensity. Note that the expected γH2Av-positive cells in the germarium in *mh[mel]* are absent under the imaging parameters used but are indeed present, see Figure S2D. (E) Ovary size estimates from *mh[mel];Chk2*^*-/-*^ and *mh[sim];Chk2*^*-/-*^ females. (F) Number of mature eggs per ovary pair from *mh[mel];Chk2*^*-/-*^ and *mh[sim];Chk2*^*-/-*^ females. In panels E and F, dotted lines correspond to *mh[mel]* and *mh[sim]* averages reported in Figure 1E and F, respectively. (*t*-test: “***” = p < 0.001, “n.s.” p > 0.05, scale bar = 25μm)

To study the cell biological basis of this block to oogenesis, we turned back to ovaries of females expressing the CRISPR-introduced *mh[mel]* or *mh[sim]* transgene under the native promoter (Figure 1B). We observed an excess of hyper-condensed nuclei in *mh[sim]* ovaries, consistent with elevated cell death (Figure 2C^28,29^). A classic trigger of cell death is the accumulation of DNA damage.^30^ To visualize DNA damage, we stained *mh[mel]* and *mh[sim]* ovaries for the double-stranded break marker, γH2Av.^31,32^ We observed elevated DNA damage signaling in *mh[sim]* ovaries (Figure 2D, S2D). This phenotype further distinguishes *mh[sim]* from *mh* null ovaries – the latter show no evidence of elevated DNA damage (Figure S2E). To address the hypothesis that MH[sim] compromises oogenesis through a DNA repair pathway, we combined *mh[sim]* with a null mutation in a DNA damage checkpoint gene. The gene, *Chk2* (also known as *mnk*), normally blocks egg production in the presence of DNA damage.^33,34^ *Chk2*^*-/-*^ ovaries bypass this checkpoint, allowing a female to make mature but damaged eggs in the presence of elevated DNA damage. We discovered that *Chk2*^*-/-*^ restores *mh[sim]* ovaries to *mh[mel]*-like ovary size and *mh[mel]*-like egg production (Figure 2E,F). However, the *mh[sim*];*Chk2*^*-/-*^ females are sterile while *mh[mel]*];*Chk*^*-/-*^ females retain fertility (Figure S2F). These data suggest that MH[sim] compromises oogenesis by triggering DNA damage.

Applying these phenotypic data to the coevolution model, we hypothesized that MH[sim]-induced DNA damage depends on the 11Mb array of 359 satellite in *D. melanogaster*. Under this model, MH[sim]-specific residues are incompatible with 359. Removing 359 should restore germline genome integrity and fertility of *mh[sim]* females. To directly test this prediction, we took advantage of a fly strain that lacks the 11Mb array of *X*-linked 359 satellite (Figure 3A^35^). We recombined this 359-deletion, called *Zygotic hybrid rescue*, or “*Zhr*,” onto both the *mh[mel]* and the *mh[sim] X* chromosomes (Figure 3A). If MH[sim]-induced toxicity depends on the presence of the 359 expansion, *mh[sim],Zhr* females should have minimal DNA damage and recover fertility. Remarkably, the 359-deletion completely restores the DNA damage marker, γH2Av, to wildtype (low) levels (Figure 3B). Consistent with restored germline genome integrity of *mh[sim]* females, we observed no difference in ovary size and no difference in egg production between *mh[mel]* and *mh[sim]* females that lack 359 (Figure 3C,D). Finally, the 359-deletion completely restores *mh[sim]* fertility to *mh[mel]* levels (Figure 3E). These data reveal that MH[sim] toxicity depends on 359, consistent with a history of coevolution between these two fast-evolving components of the Drosophila genome.

**Figure 3.**
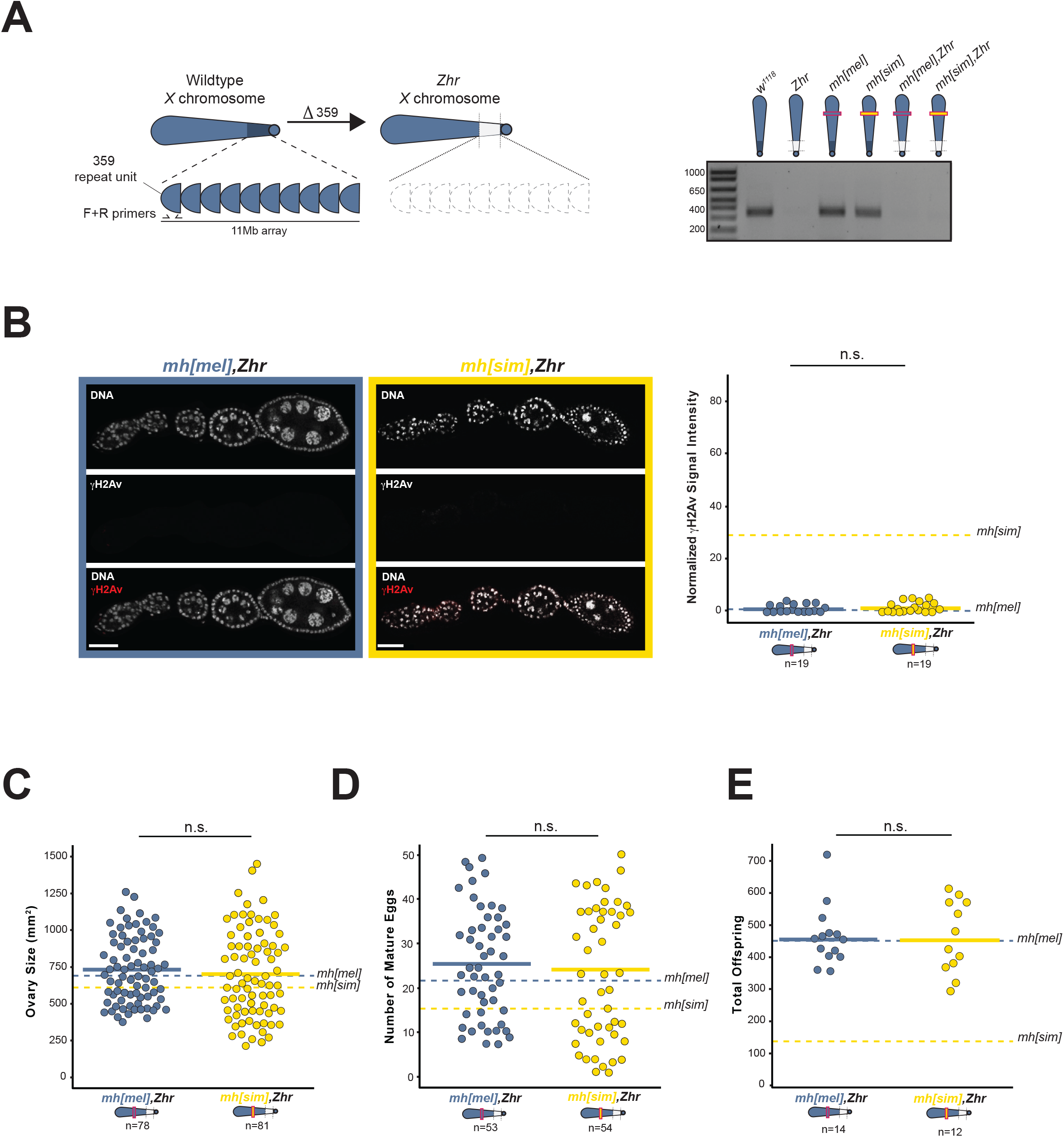
The 359 satellite deletion rescues *mh[sim]* genome integrity and fertility. (A) The *Zhr X* chromosome lacks the 11Mb pericentromeric 359 satellite array (left). A 10-cycle PCR distinguishes between wildtype 359 copy number and the 359-deletion (*Zhr*) and validates the recombined *mh[mel],Zhr* and recombined *mh[sim],Zhr X* chromosomes (right). (B) γH2Av signal in *mh[mel],Zhr* and *mh[sim],Zhr* ovaries and the quantification of normalized fluorescent signal intensity. (C) Ovary size of *mh[mel],Zhr* and *mh[sim],Zhr* females. (D) Number of mature eggs per ovary pair from *mh[mel],Zhr* and *mh[sim],Zhr* females. (E) Progeny counts from *mh[mel],Zhr* and *mh[sim],Zhr* females crossed to wildtype (*w*^*1118*^) males. In panel B, dotted lines correspond to *mh[mel]* and *mh[sim]* averages reported in Figure 2D. In panels C, D, and E, dotted lines correspond to *mh[mel]* and *mh[sim]* averages reported in Figure 1E, F, and C, respectively. (*t*-test: “n.s.” p > 0.05, scale bar = 25μm)

The observed 359-dependent toxicity rather than loss of function suggests that MH[sim] may *interfere* with the preservation of 359 integrity. To define a molecular basis for this interference, we used two, well-characterized MH homologs as guides. The MH homologs in worm (DVC-1) and human (Spartan) use the conserved Spartan metalloprotease domain to cleave DNA-protein crosslinks that form between DNA and Topoisomerase II (Top2^36–38^). These crosslinks dock Top2 at DNA entanglements, which Top2 resolves by inducing double strand breaks followed by re-ligation.^39–41^ In *D. melanogaster*, Top2 specifically cleaves 359^42^ and resolves DNA entanglements involving 359 during female meiosis.^43^ We hypothesized that MH[sim] interferes with Top2 resolution of 359 entanglements. This interference model predicts that Top2 is limiting in the presence of MH[sim]. To test this prediction, we reduced Top2 using a heterozygous loss of function mutant and overexpressed Top2 in the ovary using the UAS/ GAL4 system in an *mh[mel]* or *mh[sim]* background. Reduction of Top2 exacerbates *mh[sim]*-dependent subfertility (Figure 4A) while Top2 overexpression in the ovary completely rescues *mh[sim]* fertility (Figure 4B). The rescued *mh[sim]* ovaries also showed restored genome integrity (Figure 4C). Excess Top2 appears to overcome MH[sim] interference. Combined with the literature on DNA-Top2 crosslink cleavage by MH homologs DVC-1 and Spartan, these data raise the possibility that MH[sim] over-actively clears the 359-Top2 crosslinks necessary to resolve 359 entanglements in the female germline. Persistent DNA entanglements would trigger the observed DNA damage that blocks oogenesis progression (Figure 4D).

**Figure 4.**
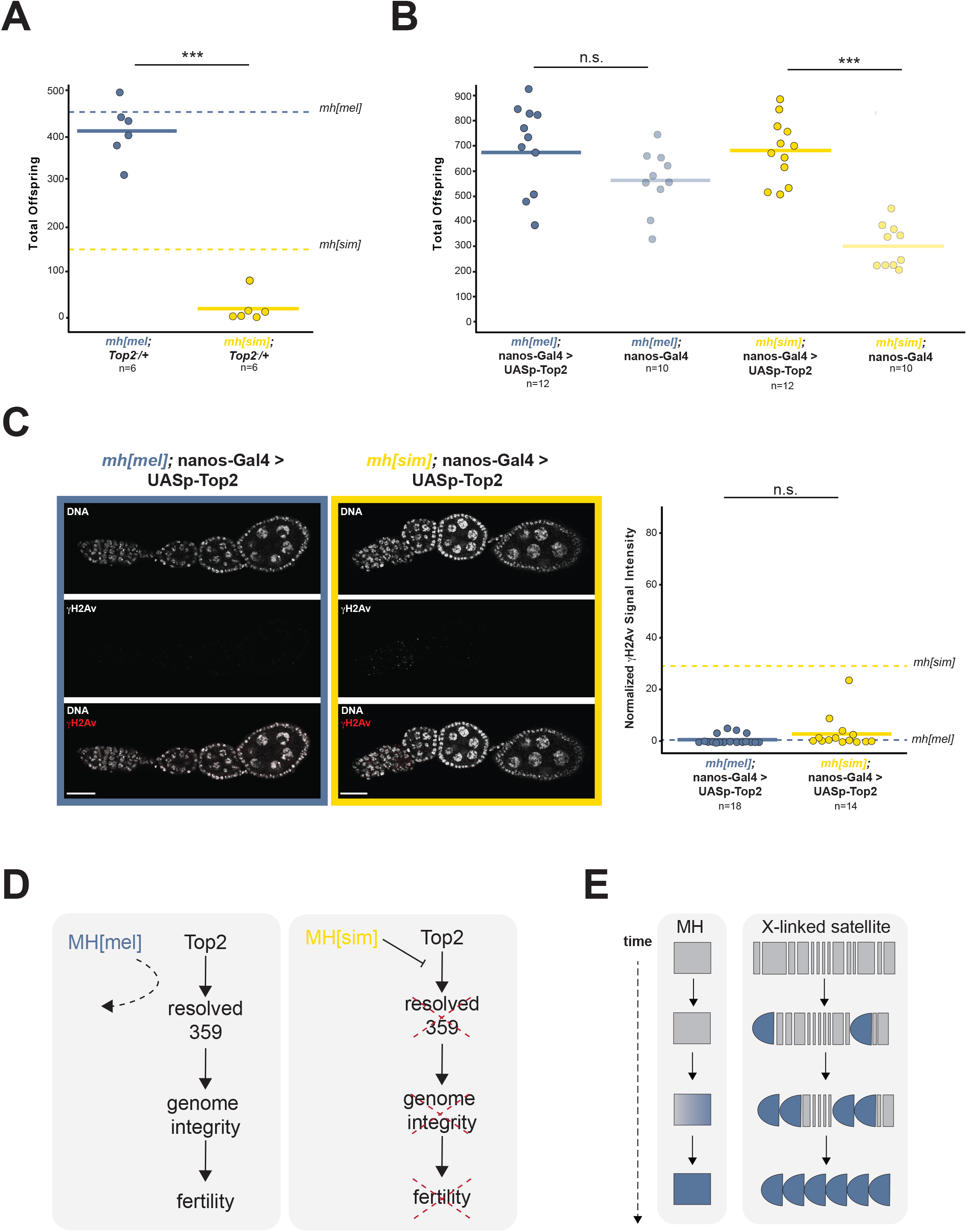
MH[sim] interferes with Top2 processing of 359 entanglements. (A) Progeny counts from *mh[mel];Top2*^−^/+ and *mh[sim];Top2*^−^/+ females crossed to wildtype (*w*^*1118*^) males. (B) Progeny counts from nos-Gal4-VP16 (female germline GAL4) driven *mh[mel];*UAS*-Top2* or *mh[sim];*UAS*-Top2* females crossed to wildtype (*w*^*1118*^) males. (C) γH2Av signal from ovaries of nos-Gal4-VP16 driven *mh[mel];*UAS*-Top2* or *mh[sim];*UAS*-Top2* females and quantification of normalized fluorescent signal intensity. (D) Model of MH[sim] interference with Top2 processing of 359 entanglements. These entanglements threaten genome integrity and ultimately, fertility. MH[mel], in contrast, has no measurable function in the ovaries, suggesting that it avoids interfering with 359 processing by Top2. (E) Model of MH evolution tracking 359 satellite proliferation. In panel A, dotted lines correspond to *mh[mel]* and *mh[sim]* averages reported in Figure 1C. In panel C, dotted lines correspond to *mh[mel]* and *mh[sim]* averages reported in Figure 2D. (*t*-test: “***” = p < 0.001, “n.s.” p > 0.05, scale bar = 25μm)

Our model is motivated in part by the observation that repeat-rich heterochromatin, but especially the 11Mb array of 359, is uniquely vulnerable to DNA entanglements.^16,43,44^ If 359 is so deleterious, how could it have proliferated? DNA satellites can behave selfishly, gaining a transmission advantage from one generation to the next.^45,46^ We suspect that such non-Mendelian segregation led to 359 proliferation, triggering MH to evolve adaptively (Figure 4E). However, we cannot formally rule out the possibility that a selection pressure distinct from 359 proliferation triggered MH adaptive evolution. Under this alternative model, the *D. melanogaster* version of MH evolved first, releasing constraint on 359 copy number. Most likely, both selection and loss of constraint operate cyclically.

Regardless of the force(s) that promoted 359 proliferation, the 359:MH system offers an important elaboration of the canonical model of intra-genomic coevolution.^10,47–49^ This canonical model posits that chromosomal proteins evolve adaptively to recognize and process novel satellite repeat variants. Under this model, the mismatched *mh[sim]* allele should fail to perform an *mh* function; that is, act as a loss of function allele. Instead, we demonstrate that *mh[sim]* is toxic, suggesting that *mh[mel]* instead evolved adaptively to *avoid interfering* with 359 processing, and likely 359 processing by Top2. The observed colocalization of MH[mel] foci with Top2 and 359 in the embryo (^24^, many cycles after the first mitosis) motivates future work dissecting the functional consequences of MH adaptive evolution during this distinct developmental stage.

359-mediated toxicity to oogenesis highlights the catastrophic functional consequences of DNA satellite evolution. Importantly, 359-mediated toxicity is also apparent in *D. melanogaster-D. simulans* hybrid embryos: a distinct, unmapped gene on *D. simulans* chromosome *2*^50–53^ interacts deleteriously with 359 to cause embryonic chromosome mis-segregation, genome instability, and lethality.^16,35,54^ This interspecies hybrid dysfunction in the embryo, together with the 359:*mh[sim]* toxicity in the ovary reported here, suggests that recurrent bouts of coevolution not only shape essential genome functions within species but also can trigger hybrid incompatibilities between species.

## ACKNOWLEDGEMENTS

We thank Isabella Farkas and Courtney Christopher for technical assistance. We also thank the Levine Lab, M. Patel, N. Phadnis, A. Das, and D. Dudka for feedback on the manuscript and the Levine Lab, P. Geyer, H. Malik, R.S. Hawley, and M. Buszczak for discussions about the project. This work was supported by a Life Sciences Research Foundation fellowship to C.L.B. and National Institutes of Health (NIH) NIGMS grant R35GM124684 to M.T.L.

## AUTHOR CONTRIBUTIONS

Conceptualization, M.T.L and C.L.B.; Methodology, M.T.L and C.L.B.; Investigation, M.T.L and C.L.B; Writing M.T.L and C.L.B; Funding Acquisition, M.T.L and C.L.B.

## STAR METHODS

### Key resources table

| REAGENT or RESOURCE | SOURCE | IDENTIFIER |
| --- | --- | --- |
| **Antibodies** |  |  |
| Mouse Monoclonal anti-FLAG M2 | Sigma-Aldrich | Cat#F3165 |
| Mouse Monoclonal 12G10 anti-αTubulin | Developmental Studies Hybridoma Bank | Cat#12G10 anti-alpha-tubulin, RRID:AB_1157911 |
| Mouse Monoclonal anti-Histone 2A Gamma Variant, Phosphorylated | Developmental Studies Hybridoma Bank | Cat#UNC93-5.2.1, RRID:AB_261807 |
| Chicken Polyconal anti-Green Fluorescent Protein | Aves Labs | Cat# GFP-1010, RRID:AB_230731 |
| **Chemicals, peptides, and recombinant proteins** |  |  |
| Phusion High-Fidelity DNA Polymerase | NEB | Cat#M0530 |
| RIPA buffer | Cell Signaling Technology | Cat#9806 |
| Protease Inhibitor Cocktail | Roche | Cat#11873580001 |
| PMSF | Cell Signaling Technology | Cat#8553 |
| Benzonase | Sigma-Aldrich | Cat#E1014 |
| ProLong Gold Antifade Mountant with DAPI | ThermoFisher Scientific | Cat#P36931 |
| **Critical commercial assays** |  |  |
| Kwikquant Western Blot Detection Kit | Kindle Biosciences | Cat#R1004 |
| **Deposited data** |  |  |
| *mh* alleles from *D. simulans* | This paper | OL546458..64 |
| **Experimental models: Organisms/strains** |  |  |
| *yw*; *nos-Cas9(II-attP40)* | Best Gene, Inc | N/A |
| *yw*; *PBac[y<sup>+</sup>-attP-9A]VK00018* | Bloomington Drosophila Stock Center | BDSC:9736 |
| *Gal4::VP16-nos* | Bloomington Drosophila Stock Center | BDSC:64277 |
| *Zhr*<sup>1</sup> | Bloomington Drosophila Stock Center | BDSC:25140 |
| *mnk*<sup>Δ6</sup> | N. Phadnis | N/A |
| *Top2*<sup>176</sup> | P. Geyer | N/A |
| *w*<sup>1118</sup> | MTL laboratory | N/A |
| *mh*<sup>1</sup> | Bloomington Drosophila Stock Center | BDSC:7130 |
| *P{gcid.EGFP.cidIII}.2* | K. McKim,<sup>55</sup> | N/A |
| **Oligonucleotides** |  |  |
| Primers to amplify *mh* alleles from *D. simulans*, see Table S1 | This paper | N/A |
| Primers to screen *mh* CRISPR transformants, see Table S1 | This paper | N/A |
| Primers to detect native *mh* locus, see Table S1 | This paper | N/A |
| Primers to screen for PCR-introduced mutations in the cloned *mh* UAS constructs, see Table S1 | This paper | N/A |
| Primers to screen for 359 deletion (*Zhr*), see Table S1 | <sup>6</sup> | N/A |
| Recombinant DNA |  |  |
| Plasmid: pBFv-U6.2 | Addgene | Addgene#138400 |
| Plasmid: pBFv-U6.2B | Addgene | Addgene#138401 |
| Plasmid: 3xFLAG:*mh[mel]* and 3xFLAG:*mh[sim]* HDR plasmids | This paper | N/A |
| Plasmid: pUASp-attB | Drosophila Genomics Resource Center | DGRC#1358 |
| Software and algorithms |  |  |
| Geneious v 11.1.5 | Biomatters | https://www.geneious.com/ |
| FIJI | ImageJ2 | https://imagej.net/software/fiji/ |

### Contact for reagent and resource sharing

Further information and requests for resources and reagents should be directed to and will be fulfilled by the Lead Contact, Mia Levine.

### Population genetic and molecular evolution analyses

We conducted population genetic analysis of *mh* using multiple alleles from both *D. melanogaster* and *D. simulans*. We obtained eight *D. melanogaster mh* alleles (coordinates *X*:15472804-15475400, dmel r6.4) from lines collected in Lyon, France.^56^ We amplified seven *D. simulans mh* alleles from lines collected in Nairobi, Kenya (Accession OL546458-OL546464)^57^. Importantly, a duplication event occurred along the *D. simulans* lineage, resulting in a full-length copy of the *mh* ortholog, and a tandem partial duplicate.^58^ To specifically amplify the full length *mh* ortholog, we designed primers that anneal to unique genomic sequence (Table S1). We then prepared genomic DNA and conducted PCR amplification followed by Sanger sequencing using standard protocols. We aligned the sequences in Geneious using the Geneious Alignment algorithm with default settings (Geneious v11.1.5, Biomatters, Auckland, New Zealand) and confirmed alignment quality by eye. We performed a McDonald-Kreitman test^26^ with the *D. melanogaster* and *D. simulans mh* coding sequences.

### Fly stock construction and husbandry

#### Constructing gene swaps

We used CRISPR/Cas9 to generate *D. melanogaster* flies that encode a transgenic *D. melanogaster* allele or a *D. simulans* allele of *mh,* integrated into the native location. We first generated a U6 promoter-driven guide RNA construct by cloning sgRNAs flanking the coding sequence of *mh* (5’: GGATTGGCCCAGGATCAACA, 3’: CGTGGAGAGCTTCTGCCGCG) into pBFv-U6.2 and pBFv-U6.2B backbones. We shuttled the 3’ sgRNA into pBFv-U6.2 to create a dual sgRNA vector (University of Utah Mutagenesis Core). In parallel, we constructed homology directed repair (HDR) plasmids encoding one kilobase homology arms 5’ and 3’ of their respective guide RNAs. Between the homology arms we synthesized a codon-optimized (for *D. melanogaster*) *mh* coding sequence from either *D. melanogaster* or *D. simulans* (GenScript, Piscataway, NJ). We N-terminally tagged each sequence with 3xFLAG along with a linker sequence (GGTGGTTCATCA). We injected the dual sgRNA vector and a single HDR plasmid into the Cas9-expressing line, *yw; nos-Cas9(II-attP40)* (BestGene Inc, Chino Hills, CA).

We crossed single males, injected as embryos, to FM7 (*X*-chromosome balancer) females. We screened F_1_ females to identify positive transformants using forward primer 5’-AAGTGTCGCGCTATTTCACC-3’ and reverse primer 5’-TCACCGTCATGGTCTTTGTAGTCCAT-3’. We then backcrossed the positive F_1_ females to FM7 males and self-crossed the balanced F_2_ progeny to generate lines homozygous for either *mh[mel]* or *mh[sim]* allele. To confirm that the introduced alleles encoded the expected sequence and in the expected location, we amplified the entire region from homozygous flies using primers that anneal outside of the homology arms (5’-AATGGATTTCGGCAAATGAG-3’, 5’-GTCGTTGTAGGAGCCCATGT-3’) and then sequenced across the entire region. We also designed primers that amplified the native *mh* locus (5’-GGCCCTGCTCATATCGTATC-3’, 5’-AAGAACCTTACTGCGTGCAAC-3’) to confirm that our final genotypes were true replacements. Primers can be found in Table S1.

#### Constructing UAS-mh and UAS-Top2 lines

We used the *ϕC31* integrase-mediated transgenesis system to introduce into the same landing site *mh* from *D. melanogaster* or *D. simulans* downstream of an “upstream activating sequence” or “UAS”^59^. Using the HDR plasmids as a template (see above), we PCR-amplified the 3xFLAG-tagged *mh* coding sequence (either *D. melanogaster* or *D. simulans)* using Phusion High-Fidelity DNA Polymerase (NEB, Ipswich, MA). We cloned the resulting PCR products into *NotI/XbaI* sites of the pUASp-attB vector (*Drosophila* Genomics Resource Center, Bloomington, IN). We confirmed the absence of PCR-introduced mutations in the cloned UAS-*mh[mel]* and UAS-*mh[sim]* alleles by direct Sanger sequencing of the constructs (Table S1). We introduced the constructs into *D. melanogaster yw; PBac[y*^+^-*attP-9A]VK00018* flies, which have an attP transgene landing site at cytological position 75A10 on chromosome *3L* (BestGene Inc, Chino Hills, CA). We next made each transgene homozygous. To overexpress the transgenic alleles, we crossed these stocks to *Gal4::VP16-nos* (BDSC #64277), which drives germline expression of transgenes downstream of UAS.

Similarly, we used the *ϕC31* integrase-mediated transgenesis system to introduce *Top2* from *D. melanogaster* downstream of an UAS promoter.^59^ We synthesized a codon-optimized (for *D. melanogaster*) *Top2* coding sequence from *D. melanogaster* (Twist, South San Francisco, CA). We N-terminally tagged each sequence with 3xHA along with a linker sequence (GGTGGTTCATCA). We introduced the constructs into *D. melanogaster yw; PBac[y*^+^-*attP-9A]VK00018* flies (see above, BestGene Inc, Chino Hills, CA). We next constructed either *mh[mel]*; UAS-*Top2[mel]* or *mh[sim]*; UAS-*Top2[mel]* stocks using balancer chromosomes. To overexpress the transgenic *Top2* allele, we crossed either *mh[mel]*; UAS-*Top2[mel]* or *mh[sim]*; UAS-*Top2[mel]* males to either *mh[mel]*; *Gal4::VP16-nos* or *mh[sim]*; *Gal4::VP16-nos* females, respectively.

#### Zhr rescue stocks

To generate stocks that encode both the *X-*linked *mh*-transgene and the *X-*linked 359 satellite deletion (*Zhr*^*1*^, BDSC #25140), we first generated trans-heterozygote females. We crossed these trans-heterozygote females to FM7 males and used PCR to assay individual recombinant male progeny for the presence of both the *mh* transgene and *Zhr.* We detected the *mh* transgenes with forward primer 5’-AAGTGTCGCGCTATTTCACC-3’ and reverse primer 5’-TCACCGTCATGGTCTTTGTAGTCCAT-3’. To detect the *Zhr* mutation (*i.e.*, 359 satellite deletion), we used forward primer 5’-TATTCTTACATCTATGTGACC-3’ and reverse primer 5’-GTTTTGAGCAGCTAATTACC-3’.^6^ Performing a 10-cycle PCR at an annealing temperature of 52C yields a band only in the presence of the 11Mb 359 satellite array (Figure 3A). We backcrossed males positive for both the *mh* transgene and *Zhr* mutation to FM7 females to generate a permanent stock.

#### Additional stocks

Heterozygote *mh[mel]/mh[sim]* females were generated by crossing *mh[mel]* females to *mh[sim]* males.

We used a +/FM7; +/CyO stock to generate flies encoding both the *mh* transgene at the native locus (chromosome *X*) and the *Chk*^-/-^ (*mnk*) mutation (chromosome *2*). The *mnk*^*l6*^ stock^34^ was a gift from N. Phadnis.

To generate heterozygous *Top2* hypomorph females, we also used a +/FM7; +/CyO stock to generate flies encoding both the *mh* transgene at the native locus (chromosome *X*) and a heterozygous *Top2*^*176*^/CyO mutation (chromosome *2*). The *Top2*^*176*^ stock^60^ was a gift from P. Geyer.

### Immunoblotting

To assay 3x-FLAG MH protein abundance in the ovary, we dissected 20 ovary pairs in 1X PBS and ground the material in RIPA buffer (Cell Signaling Technology, Danvers, MA), Protease Inhibitor Cocktail (Roche, Basel, Switzerland), and 2X PMSF (Cell Signaling Technology, Danvers, MA). To promote solubility, we incubated the lysate in benzonase (Sigma Aldrich, St. Louis, MO) for 1hr at 4C. We used 20µg of lysate and probed with 1:10,000 anti-FLAG (M2, Sigma Aldrich, St. Louis, MO) or 1:1000 anti-αTubulin (Developmental Studies Hybridoma Bank, Iowa City, IA) and 1:1000 anti-mouse HRP secondaries (Kindle Biosciences, Greenwich, CT). We exposed blots with Kwikquant Western Blot detection kit and imaged with a Kwikquant imager (Kindle Biosciences, Greenwich, CT).

### Fertility assays

#### Female fertility

To assay female fertility, we first aged virgin females 3-5 days. For each replicate vial, we crossed four virgin females to four *w*^*1118*^ males. We conducted all crosses on molasses food at 24C. We flipped the parents onto new food every three days over the course of nine days and counted all progeny that emerged.

#### Ovary size and mature egg counts

To determine the number of mature eggs and ovary size from focal genotypes, we first dissected ovary pairs in 1X PBS and imaged at 8X magnification with a Leica DFC7000 T camera. We quantified the area of each ovary using the polygon tool in FIJI^61^ to define the borders of the tissue. For each ovary pair, we used the Freehand tool in FIJI to calculate the area (mm^2^) within these boundaries for each individual ovary. After imaging, we counted the number of eggs that contain elongated dorsal appendages (stages 13 and 14).

### Immunofluorescence

We conducted immunofluorescence on ovaries following the protocol described in.^62^ We stained ovaries with anti-FLAG (1:3000, M2, Sigma Aldrich, St. Louis, MO) and anti-γH2Av (1:1000, a gift from R. S. Hawley). We mounted ovaries with ProLong Gold Antifade Reagent with DAPI (Thermo Fisher Scientific, Waltham, MA). We imaged slides at 63X magnification on a Leica TCS SP8 Four Channel Spectral Confocal System. For each experiment, we used the same imaging parameters across genotypes.

We conducted immunofluorescence on embryos collected in a 0-70 minute window from *mh[mel]*, *mh[sim]*, or *mh*^*1*^ females crossed to males homozygous for *P{gcid.EGFP.cid}III.2* (^55^, a gift from K. McKim). We followed the protocol described in^63^ to fix and stain the embryos with anti-GFP (1:1000, Aves Labs, Tigard, OR). We mounted and imaged the embryos as described above.

### Analysis of cytological data

#### Cell death quantification

To quantify the incidence of cell death, we mounted fixed whole ovaries with ProLong Gold Antifade Reagent with DAPI (Thermo Fisher Scientific, Waltham, MA) and imaged at 63X magnification on a Leica TCS SP8 Four Channel Spectral Confocal System using the tile scanning and merging feature. We identified the number of ovarioles that contained egg chambers with >1 condensed, signal-saturated nurse cell nuclei. We then divided this number by the total number of ovarioles present in each ovary to determine the fraction of cell death incidence in *mh[mel]* and *mh[sim]* ovaries.

#### Immunofluorescence quantification

To quantify the average fluorescence of γH2Av in *mh[mel]* and *mh[sim]* ovaries we outlined a representative stage four egg chamber with the Freehand tool in FIJI^61^. We calculated the fluorescent signal intensity using the polygon tool in FIJI to define the borders of the tissue. We used the Measure tool in FIJI to calculate the mean pixels within these boundaries. We normalized the fluorescent signal intensity of *mh[mel]*, *mh[sim], and mh*^*1*^ to the mean intensity signal of the *mh[mel]*. Similarly, the fluorescent signal intensity of *mh[mel],Zhr* and *mh[sim],Zhr* Was normalized to the mean intensity signal of *mh[mel],Zhr*. Finally, the fluorescent signal intensity of *mh[mel]*; UAS-*Top2[mel]* and *mh[sim]*; UAS-*Top2[mel]* was normalized to the mean intensity signal of *mh[mel]*; UAS-*Top2[mel]*.

## SUPPLEMENTARY FIGURE LEGENDS

**Figure S1.**
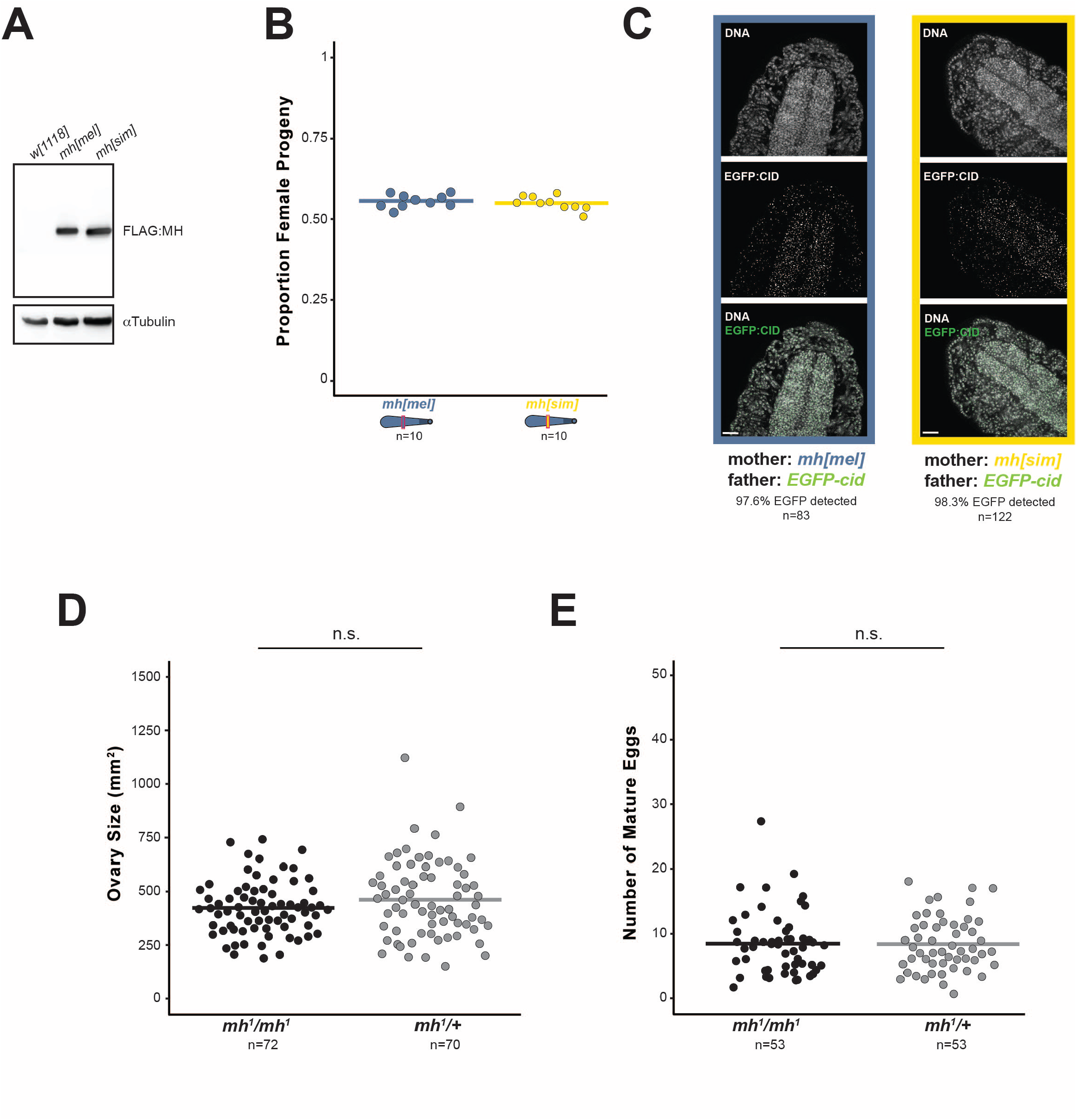
MH[sim] is expressed at comparable levels to MH[mel] and does not phenocopy *mh* null. (A) Western Blot of *mh[mel]* and *mh[sim]* ovaries probed with anti-FLAG and anti-αTubulin. (B) Proportion of female progeny from *mh[mel]* and *mh[sim]* females. (C) Representative images of diploid embryos from crosses between *EGFP-cid* fathers and *mh[mel]* or *mh[sim]* mothers stained with anti-GFP. Diploid embryos are GFP-positive and haploid embryos are GFP-negative. (D) Ovary size estimates from *mh*^*1*^ homozygous and *mh*^*1*^/*+* heterozygous females. (F) Number of mature eggs per ovary pair from *mh*^*1*^ homozygous and *mh*^*1*^/+ heterozygous females. In panel B, dotted lines correspond to *mh[mel]* and *mh[sim]* averages reported in Figure 1C. (*t*-test: “n.s.” p > 0.05, scale bar = 25μm)

**Figure S2.**
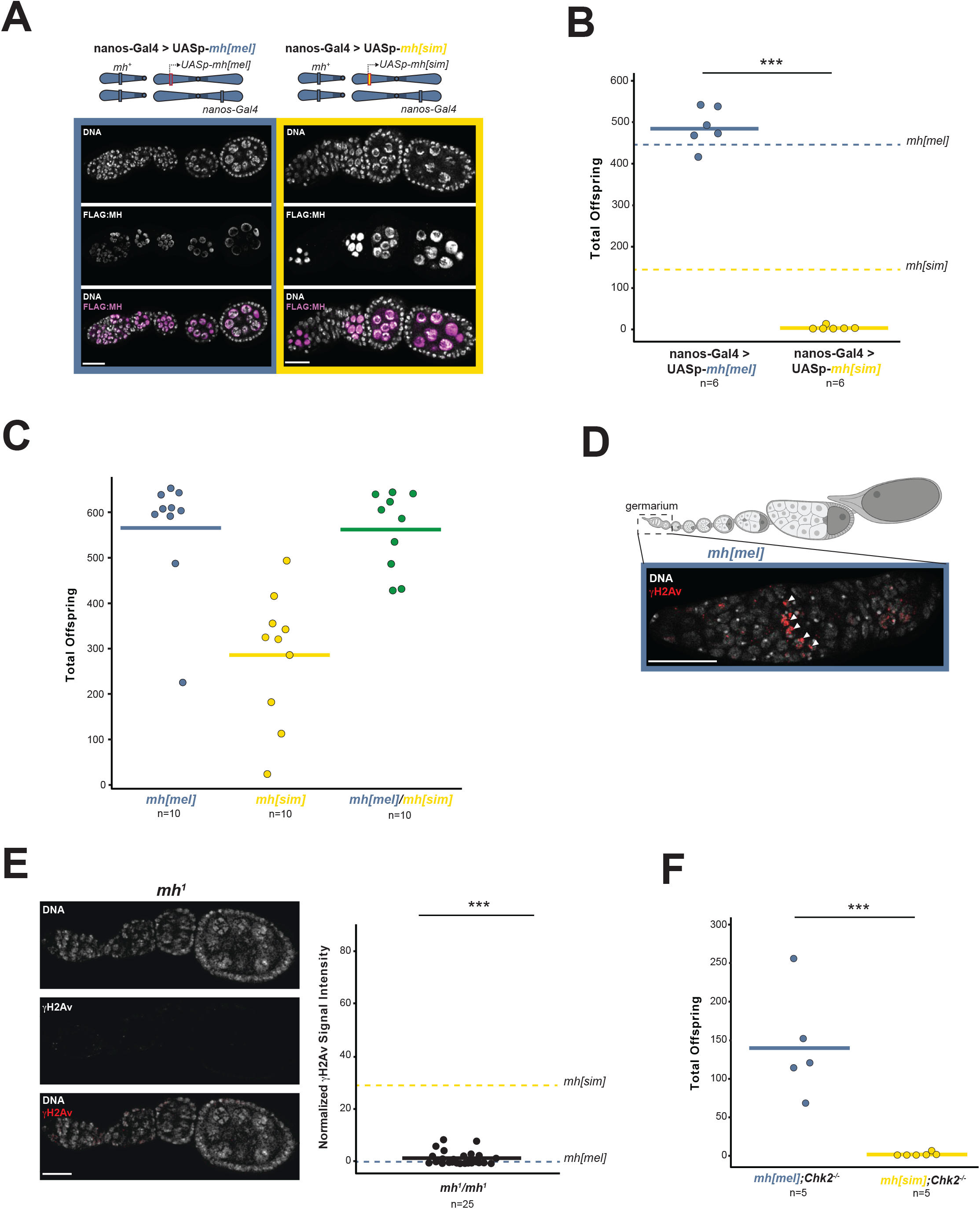
MH[sim] toxicity is dose-dependent, phenotypically distinct from *mh* null, and acts through a DNA damage pathway to block oogenesis. (A) Anti-FLAG staining of *mh[mel]* and *mh[sim]* upon overexpression under the UAS-GAL4 system. The driver nos-GAL4-VP16 expresses the *mh* transgene in the female germline only. Corresponding chromosome *X* (wildtype) and chromosome *3* are shown above images. (B) Progeny counts of the nos-Gal4-VP16 driven UASp-*mh[mel]* or UASp-*mh[sim]* females crossed to wildtype (*w*^*1118*^) males. Note that wildtype *mh* is present in these genotypes. (C) Total offspring of *mh[mel], mh[sim],* and *mh[mel]/mh[sim]* heterozygous females crossed to wildtype (*w*^*1118*^) males. (D) *Drosophila* ovariole (above) showing the germarium where meiotic recombination occurs. Germarium of an *mh[mel]* female (below) showing programmed double stranded breaks (arrowheads) occurring in “region 2A” of the germarium detected under higher laser power relative to images displayed in Figure 3B. These double stranded breaks are repaired via meiotic recombination pathways. (E) γH2Av signal in *mh*^*1*^ ovaries and quantification of normalized fluorescent signal intensity. (F) Progeny counts from *mh[mel];Chk2*^*-/-*^ and *mh[sim];Chk2*^*-/-*^ females crossed to wildtype (*w*^*1118*^) males. Note that *mh[mel];Chk2*^*-/-*^ reduced progeny counts are due to the *Chk2*^*-/-*^ mutation^64^. In panel E, dotted lines correspond to *mh[mel]* and *mh[sim]* averages reported in Figure 2D. (*t*-test: “***” = p < 0.001, “n.s.” p > 0.05, scale bar = 25μm)

## SUPPLEMENTARY TABLE LEGEND

**Table S1.**
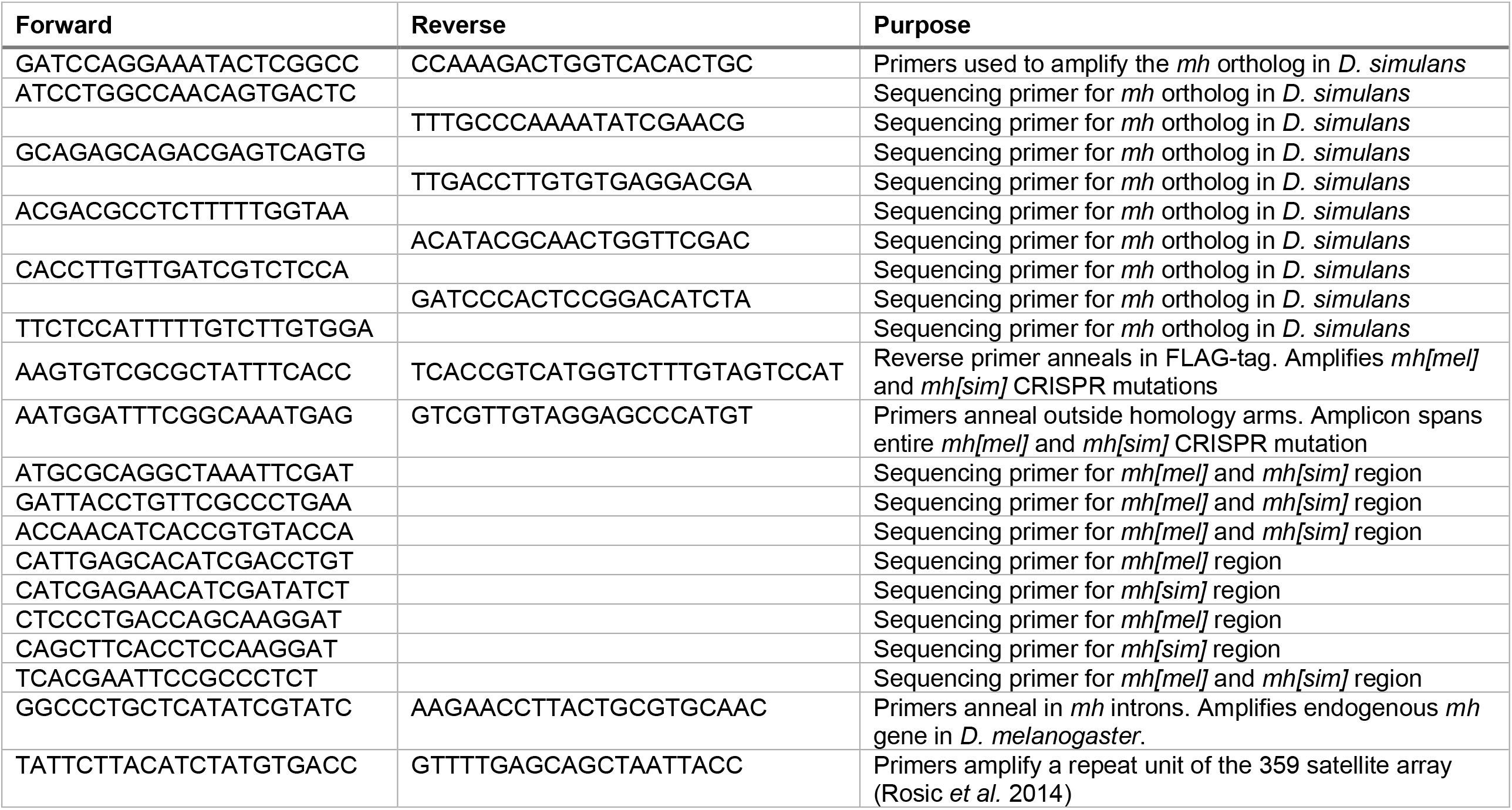
Primers used in this study.

